# CryoSCAPE: Scalable Immune Profiling Using Cryopreserved Whole Blood for Multi-omic Single Cell and Functional Assays

**DOI:** 10.1101/2024.08.20.608826

**Authors:** Alexander T. Heubeck, Cole Phalen, Neel Kaul, Peter J. Wittig, Jessica Garber, Morgan Weiss, Palak C. Genge, Zachary Thomson, Claire Gustafson, Julian Reading, Peter J. Skene

**Affiliations:** Allen Institute for Immunology, Seattle, Washington, U.S.A; Western University of Health Sciences, College of Osteopathic Medicine of the Pacific Northwest, Lebanon, Oregon, U.S.A

## Abstract

**Background:** The field of single cell technologies has rapidly advanced our comprehension of the human immune system, offering unprecedented insights into cellular heterogeneity and immune function. While cryopreserved peripheral blood mononuclear cell (PBMC) samples enable deep characterization of immune cells, challenges in clinical isolation and preservation limit their application in underserved communities with limited access to research facilities. We present CryoSCAPE (Cryopreservation for Scalable Cellular And Proteomic Exploration), a scalable method for immune studies of human PBMC with multi-omic single cell assays using direct cryopreservation of whole blood.

**Results:** Comparative analyses of matched human PBMC from cryopreserved whole blood and density gradient isolation demonstrate the efficacy of this methodology in capturing cell proportions and molecular features. The method was then optimized and verified for high sample throughput using fixed single cell RNA sequencing and liquid handling automation with a single batch of 60 cryopreserved whole blood samples. Additionally, cryopreserved whole blood was demonstrated to be compatible with functional assays, enabling this sample preservation method for clinical research.

**Conclusions:** The CryoSCAPE method, optimized for scalability and cost-effectiveness, allows for high-throughput single cell RNA sequencing and functional assays while minimizing sample handling challenges. Utilization of this method in the clinic has the potential to democratize access to single-cell assays and enhance our understanding of immune function across diverse populations.

## Background

The rapidly evolving field of single cell technologies has become a central component in advancing our understanding of the human immune system, revealing the cellular heterogeneity that underlies immune function. Analyzing protein abundance, gene expression, and epigenetics at single-cell resolution provides new observations into cell phenotypes and states in both health and disease, with the potential to identify novel biomarkers and therapeutic targets^1,2^. A critical aspect to utilizing these tools is the sourcing, isolation, and preservation of clinical samples from patients for studies performed downstream in research laboratories. Peripheral blood mononuclear cells (PBMCs) are one of the most common sample types for studying the immune system^3^ and serve as a valuable source for such studies, offering a snapshot of the immune system at the time of sample collection that can be cryopreserved for extended periods and analyzed using state of the art technologies years later. Cryopreserved PBMC samples also enable functional assays many years after collection, including cellular stimulations and genetic manipulations^4,5,6,7^.

Although isolating and cryopreserving PBMCs is the current standard method in immunology research studies, there are challenges related to PBMC isolation and preservation in the clinic that limit the scope of sample collection. In the clinical isolation of PBMCs for immune studies, the primary obstacles stem from the time and method sensitive nature of the process. Delays in sample processing of more than 4 hours after blood draw can have profound changes in the single-cell transcriptome and composition of the plasma proteome^8,9^. Standardizing sample handling methods becomes particularly challenging in clinical cohorts across multiple sites, introducing variability that may obscure important biological differences^7^. Additionally, limited access to specialized equipment and trained personnel acts as a barrier, preventing many clinical sites from participating in studies that require PBMC isolation. This limitation, especially prevalent in low-income or underserved areas, hinders access to diverse patient samples^10,11^. Commercial options that address these issues with point of collection fixation exist, but were largely designed for flow cytometry and are not compatible with most next generation sequencing assays^12,13^. Other approaches utilizing cryopreservation of whole blood have shown extension to single cell RNA sequencing but without faithfully maintaining cell proportions or demonstration of functional assays^14^. Addressing these obstacles is crucial for enabling more impactful analysis of human immunology, with the potential to expand current studies to a broader and more diverse population.

In addition to the challenges of human PBMC sample collection and preservation, the current constraints of single-cell technologies pose significant obstacles to conducting large-scale studies in many research settings. In the case of single cell sequencing, prohibitively high costs and method complexity often restrict the number of patient samples that can be feasibly analyzed and diminish the efficacy of clinical datasets^15^. Many single-cell assays also necessitate the use of live cells, thereby limiting access to in vitro time course studies that are optimally conducted using fixatives to capture cell states at precise time points^16,17^. Improving upon the cell handling and assay scalability limitations of single cell sequencing, as well as the challenges in clinical PBMC preservation, would democratize access to these important technologies, unlocking new possibilities for clinical discovery and translational impact.

To address the challenges of conventional human immune studies, we optimized an accessible sample collection methodology with a scalable and cost effective method for generating single cell sequencing datasets combined with functional assays using human PBMCs called CryoSCAPE (Cryopreservation for Scalable Cellular And Proteomic Exploration). This approach utilizes direct cryopreservation of whole blood using freezing media containing DMSO in a similar fashion to standard PBMC cryopreservation. Red blood cell and granulocyte removal that would usually be completed by density gradient PBMC isolation in the clinic are performed on the samples at the research site to reduce handling time in the clinic and enable scalable multiplexed scRNA-seq readouts. To demonstrate the utility of CryoSCAPE, we show that relative cell proportions, single cell gene expression, single cell protein expression and epigenetic profiling of cryopreserved density gradient isolated PBMCs and cryopreserved whole blood were highly comparable. The method was also optimized for high throughput and low cost sample processing through automation, as demonstrated by analysis of highly multiplexed fixed scRNA-seq results from sixty cryopreserved whole blood samples in a single batch. In addition to multi-omic single cell readouts, the developed method enables stimulation of PBMC from cryopreserved whole blood, verified through in vitro immune stimulation assays. The integration of this methodology into research and clinical settings has the potential to broaden the application of single-cell assays, especially in underrepresented communities, and enhance our understanding of immune function across diverse populations.

## Results

### Cryopreserved whole blood samples demonstrate comparable results to cryopreserved density gradient isolated PBMC using single cell multi-omics readouts

To improve accessibility to clinical sample collection of human peripheral blood mononuclear cells (PBMCs), we optimized a whole blood cryopreservation method for use in clinical settings without access to equipment required for density gradient PBMC isolation (Fig. 1a). The method uses a freezing media composed of 15% dimethyl sulfoxide (DMSO) as a cryopreservation agent, diluted with CryoStor preservation media as a serum-free protein supplement. In this method, whole blood drawn in sodium heparin tubes is combined 1:1 with 15% DMSO freezing media in 5 mL cryopreservation tubes to achieve a final DMSO concentration of 7.5%. The whole blood is thoroughly mixed in the freezing media by inverting the tubes, then transferred into a slow freezing container. Samples can be safely stored at -80°C in a freezer or in dry ice before being shipped to the research laboratory for long term storage in liquid nitrogen. This simple and immediate cryopreservation method allows for preservation of PBMCs with no delay in processing time, and without specialized technicians or equipment, as required with standard PBMC density gradient isolation methods.

**Figure 1:**
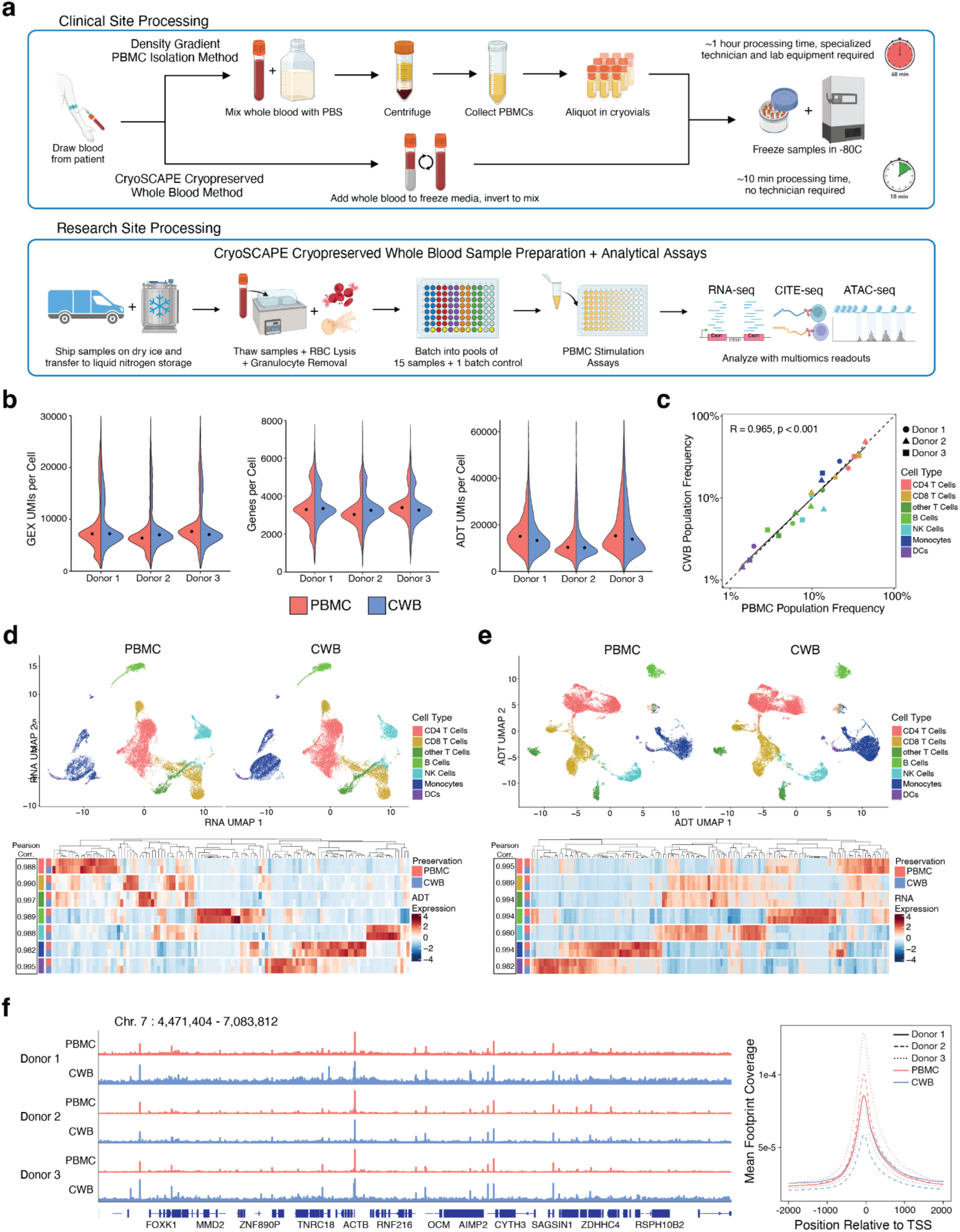
Whole blood cryopreservation in comparison to density gradient isolated PBMC cryopreservation for clinical site processing and multi-modal sequencing assays. **a**. Schematic of clinical site processing for whole blood cryopreservation versus density gradient isolated PBMC cryopreservation demonstrating reduced processing time and workflow complexity. **b.** scRNAseq data quality metrics from individual donors with density gradient isolated PBMCs (red) and cryopreserved whole blood (blue) including gene expression (GEX) unique molecular identifiers (UMIs) per cell, genes per cell, and antibody derived tag (ADT) UMIs per cell. **c.** Correlation of cell type population frequencies between donor matched cryopreserved whole blood and density gradient isolated PBMC samples (Dashed line: R=1, solid line: regression line, R=0.97, p<0.001). **d.** (Top) UMAP visualization of gene expression from cryopreserved whole blood and density gradient isolated PBMC samples overlaid with cell type labels. (Bottom) Relative gene expression levels for the highest expressing genes per cell type each cryopreservation method. High correlation is measured for each cell type indicating consistent gene detection between methods (R>0.979, p<0.001). **e.** (Top) UMAP visualization of ADT protein expression from cryopreserved whole blood and density gradient isolated PBMC samples overlaid with cell type labels. (Bottom) ADT expression levels for each cryopreservation method. High correlation is measured for each cell type indicating consistent protein detection between methods (R>0.981, p<0.001). **f.** (Left) Bulk ATAC-seq results showing a representative genome track (Chr. 7 : 4,471,404 - 7,083,812) with equivalent chromatin accessibility for cryopreserved whole blood and density gradient isolated PBMC samples across 3 donors. (Right) Distribution of footprints and position relative to TSS: total coverage of Tn5 footprints summed across all transcription start sites (TSS). Tn5 footprints are 29 bp regions comprising the 9 bp target-site duplication (TSD) and 10 bp on either side, which represent accessible chromatin for each transposition event.

By simplifying the clinical site requirements for sample cryopreservation, additional processing at the research site is required to fully enrich PBMCs for single cell assays. Once the cryopreserved whole samples have been thawed in a 37°C water bath, red blood cells (RBCs) are removed using a RBC lysis buffer, as they can reduce gene detection in scRNA-seq readouts^18^. Granulocytes from cryopreserved whole blood samples typically have very low viability upon thaw, and can cause cell clumping through release of DNA and lysosomal enzymes upon thawing^13,18^. Samples are incubated with DNase I solution to reduce clumping prior to further enrichment^19^. Granulocytes, dead cells, and other debris that are removed with density gradient PBMC isolation must also be removed from the samples, typically using a cell isolation method such as fluorescence-activated cell sorting (FACS). While this additional enrichment increases the workload at the research site, removing that responsibility from the clinical side enables more sample collection centers to participate in immunological studies.

This whole blood cryopreservation technique was evaluated for compatibility with multi-omic sequencing assays through a comparative analysis of PBMCs preserved using the cryopreserved whole blood method versus the conventional density gradient PBMC isolation approach (Fig. 1a). Whole blood samples were obtained from three donors and subjected to cryopreservation using both the whole blood and density gradient PBMC cryopreservation methods, with subsequent storage for a minimum of 6 months. Analysis of PBMCs from each method included enrichment with FACS (Supplement Fig. 1) followed by deep molecular characterization via scRNA-seq, scCITE-seq, and bulk ATAC-seq, to evaluate differences in gene expression, surface protein expression, and chromatin accessibility, respectively. Gene expression results from healthy donors (n=3) were captured using the 10x Genomics Flex single cell fixed RNA-seq kit, resulting in data from 21,023 and 17,780 single cells generated from cryopreserved whole blood and density gradient isolated PBMCs, respectively. Protein expression was measured using CITE-seq antibody derived tags (ADTs) in conjunction with the 10x Flex single cell fixed kit, for matched gene and protein results from the same single cells. Both the cryopreserved whole blood and density gradient isolated PBMC methods produced high quality data with comparable unique gene expression molecular identifiers (UMIs) per cell, median gene counts per cell, and antibody derived tags (ADTs) UMIs per cell (Fig 1b). After removal of doublets and normalization, samples were labeled using Seurat label transfer to identify 7 major immune cell types (CD4 T Cells, CD8 T Cells, Other T Cells, B Cells, NK Cells, Monocytes, and Dendritic Cells (DCs)) based on gene expression (Fig. 1c). Relative population frequencies of the major cell types displayed high correlation between cryopreserved methods (R=0.970, p<0.001), indicating no reduction of specific cell populations in cryopreserved whole blood.

Similarity between cryopreservation methods is further confirmed with combined donor Uniform Manifold Approximation Projections (UMAPs) clustered on gene expression, showing high similarity of cell type clustering (Fig 1d). To assess cell type specific differences, the average expression of the highest expressing genes per cell type were compared across cryopreservation methods (142 total genes). Gene expression was highly correlated for all cell types (R>0.979), again highlighting the similarity of gene expression between cryopreserved whole blood and density gradient isolated PBMC. Protein expression UMAPs were also labeled using the same major cell type label transfers, demonstrating a highly similar surface protein profile between the cryopreservation methods (Fig. 1e). Cell type specific protein differences were evaluated by comparing the average expression of ADTs with expression levels above isotype controls (116 total proteins). Similar to the gene expression data, protein expression was highly correlated for all cell types (R>0.981), further demonstrating the viability of using cryopreserved whole blood for single cell sequencing readouts with no loss of resolution compared to standard PBMC cryopreservation methods.

In addition to gene and protein expression, epigenetic profiling is an important approach to measure enhancer and transcription factor accessibility but has also been shown to be sensitive to delays in PBMC processing^17^. Chromatin accessibility was evaluated using ATAC-seq on bulk cell populations in both whole blood and density gradient isolated PBMC cryopreservation methods. The profiles of cryopreserved whole blood and density gradient isolated PBMCs show similar profiles with enrichment around transcriptional start sites (TSS) at a representative region of the genome (Fig. 1f, Supplement Table 2). To evaluate this on a genome wide scale, all libraries were equally downsampled to 14 million total mapped sequenced reads and total coverage of Tn5 footprints summed across all TSS. Clear enrichment across TSS regions was observed in both sample cryopreservation methods. Reduced peak height at the TSS in cryopreserved whole blood samples was observed, with the ratio of the maximum peak height near the TSS between cryopreserved whole blood and density gradient isolated PBMCs varying between samples (0.99, 0.56 and 0.59 in Donors 1, 2 and 3, respectively). This decreased signal to noise ratio in the cryopreserved whole blood may be due to extracellular debris generated from the whole blood cryopreservation workflow, but still allows for interpretation of chromatin accessibility. The successful interrogation of cryopreserved whole blood samples using these multi-omic assays enables CryoSCAPE to be used for simpler sample collection for immunological studies.

### Scaling cryopreserved whole blood sample throughput using 10x Flex scRNA-seq technology and automation

Analysis of circulating PBMC using scRNA-seq is a powerful tool to gain insight into the human transcriptome; however, the cost and scalability of these assays still remains a significant obstacle to more widespread adoption. Droplet based single cell sequencing technologies that utilize a poly-A based mRNA capture method for full transcriptome coverage have improved single cell throughput in recent years, but have yet to lower the assay cost enough to enable large scale studies^20^. A recently released method for measuring gene expression from mRNA of fixed cells, the Chromium Single Cell Gene Expression Flex kit from 10x Genomics, employs a paraformaldehyde cell fixation step followed by the hybridization of a barcoded probe panel to capture transcriptional activity. With the barcoded probe panel utilized in 10x Flex scRNA-seq, doublets can be resolved in-silico, permitting a higher cell loading concentration and assay throughput, significantly driving down the cost per cell. Further cost efficiencies can be found with workflow automation allowing for the analysis of batches of 64 or more samples, providing an 6-fold cost reduction per sample ($526 Flex vs. $3,604 3’ v3.1, Supplement Table 3). In addition to the cost advantages due to multiplexing, the 10x Flex scRNA-seq method also displays improved sensitivity and is more compatible with high throughput processing by incorporating additional stop points and a less complex overall workflow.

To evaluate the 10x Flex scRNA-seq as a higher throughput and cost effective alternative to 10x 3’ v3.1 scRNA-seq, gene expression profiles generated from each method were compared using density gradient isolated PBMCs from healthy donors (n=3). The 10x Flex scRNA-seq kit showed improved assay sensitivity, detecting ∼70% more genes and equal numbers of unique molecular identifiers (UMIs) from 75% of captured reads compared to the 10x 3’ v3.1 kit despite libraries being sequenced at ∼75% read depth (Fig. 2a). The 10x Flex method also shows improved detection of genes per cell (2746 ± 14 s.e.m.) over 10x 3’ v3.1 (1527 ± 19 s.e.m.) (Fig. 2b). Despite the differences in gene detection efficiency, concordant cell population frequencies were observed based on cell label transfer between 10x Flex and 10x 3’ v3.1 (R=0.995, pval<0.001), suggesting that there was no observable effect of probe hybridization bias towards a specific cell type leading to differences in cell type labeling across 3 donors (Fig. 2c).

**Figure 2:**
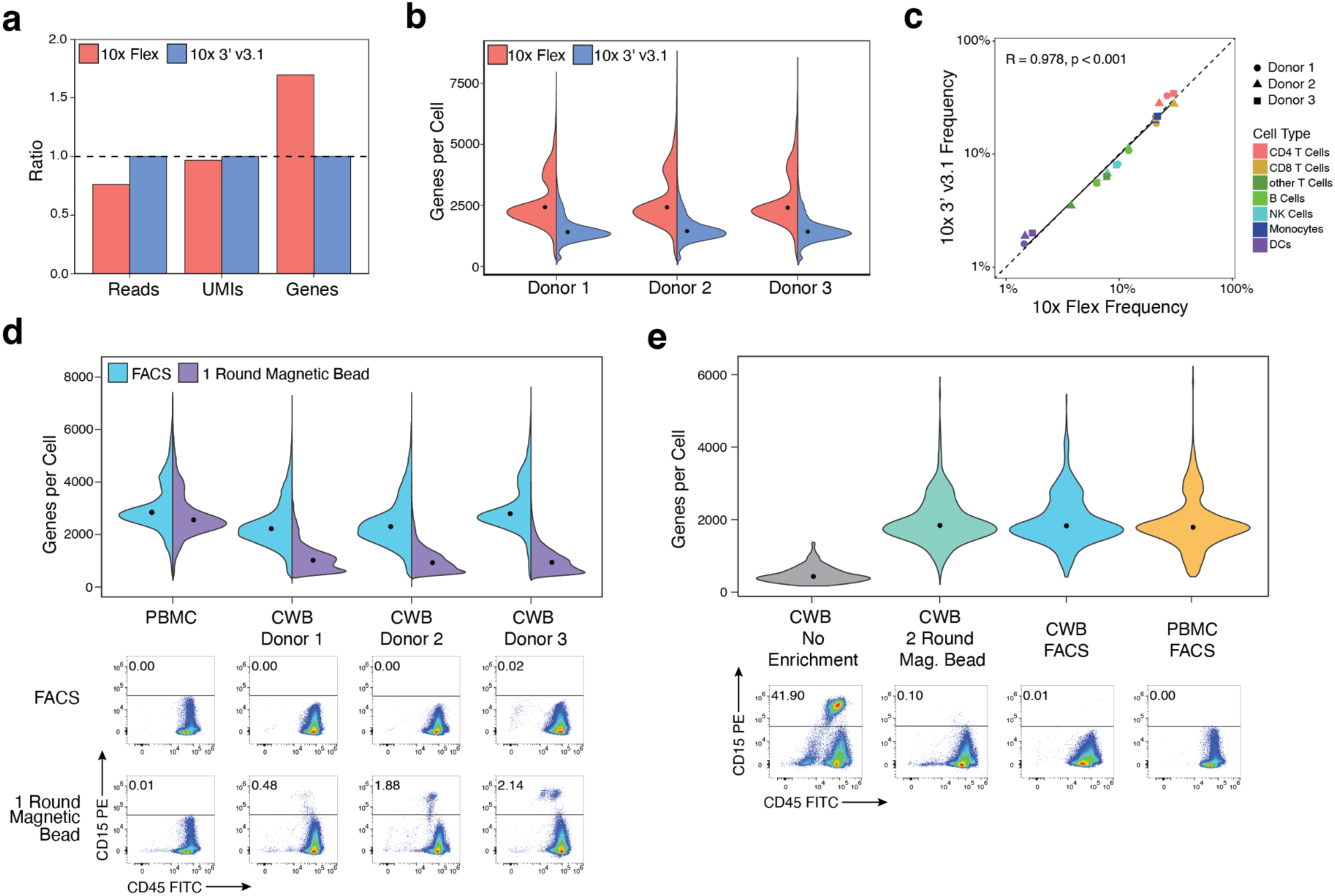
Development of 10x Flex multiplexed scRNA-seq methodology as a high throughput method for single cell transcriptome analysis of cryopreserved whole blood. **a.** Ratio of 10x Flex (red) to 10x 3’ v3.1 (blue) quality metrics comparing number of reads, UMIs, and genes detected in scRNAseq libraries prepared from PBMC donors (n=3). **b.** Genes per cell recovered from individual donors (n=3) using 10x Flex and 10x 3’ v3.1 methods. **c.** Correlation of cell type population frequencies between donor matched samples analyzed using 10x Flex and 10 3’ v3.1 methods (Dashed line equivalent to R=1, solid line equivalent to regression line, R=0.995, p<0.001). **d.** (Top) Genes per cell recovered using the 10x Flex kit with cryopreserved whole blood and density gradient isolated PBMC samples enriched using a single round of magnetic bead based enrichment (purple) versus FACS (blue). (Bottom) Remaining granulocyte contamination after enrichment by flow cytometry analysis. Relatively low numbers of granulocytes impact recovered genes per cell. **e.** (Top) Genes per cell recovered using the 10x Flex kit with cryopreserved whole blood samples with no enrichment (gray), improved 2 round magnetic bead enrichment (green), and FACS (blue). Improved magnetic bead enrichment recover similar genes per cell compared to density gradient isolated PBMCs analyzed using the same kit (yellow). (Bottom) Remaining granulocyte contamination after enrichment by flow cytometry analysis. Improved magnetic bead enrichment removes granulocytes to a sufficient level.

The performance of the 10x Flex scRNAseq kit was evaluated in conjunction with the developed whole blood cryopreservation method to enable a low cost-scalable method for high throughput single cell transcriptome analysis. While FACS is a highly effective method for isolating high purity cells for scRNA-seq, sorting cryopreserved whole blood samples can take up to 15 minutes per sample (24 hours for 96 samples) due to high debris content, severely limiting the throughput for larger batches of samples. Negative selection with magnetic bead enrichment, where non-target cells and debris are bound with magnetic beads and removed with a strong magnet, is an alternative purification method that can be easily scaled. Magnetic bead enrichment and FACS methods were compared for isolating PBMCs from cryopreserved whole blood (n=3) and density gradient isolated PBMCs (n=1), measured by 10x Flex scRNA-seq quality metrics and enrichment of PBMCs using flow cytometry analysis (Fig. 2d). As expected, FACS enrichment of viable leukocytes produced comparable UMIs and genes per cell between samples isolated from cryopreserved whole blood and density gradient isolated PBMCs. The initial magnetic bead enrichment method was tested using a single round of bead binding of a human dead cell removal kit to remove the low viability granulocyte compartment of the cryopreserved whole blood. Samples enriched using this magnetic bead depletion showed high quality metrics using the 10x 3’ v3.1 kit (Supplement Fig. 2). However, despite enriching the PBMCs from cryopreserved whole blood to at least 95% purity, a significant loss in 10x Flex library quality was observed, as less than half of the genes per cell were detected compared to cells enriched from density isolated PBMCs using the same method (Fig. 2d).

The magnetic bead enrichment method was improved by adding PE conjugated antibodies targeting CD15 and CD66b followed by binding of anti-PE magnetic beads in addition to the dead cell removal beads to target any remaining granulocytes in the samples. This step was then repeated for a total of two rounds to further isolate the target cells to greater than 99% purity, which closely matched the purity of FACS sorted cells (Fig. 2e). This additional removal of granulocytes improved the genes per cell detected in the magnetic bead depleted samples to the same level as the FACS sorted or density gradient isolated PBMC samples, demonstrating the utility of this scalable sample enrichment method with the 10x Flex scRNAseq assay (Fig. 2e). A cryopreserved whole blood sample with no enrichment included as a control produced extremely low detection of genes per cell, further highlighting the need to completely remove all granulocytes from a sample before using the 10x Flex scRNA-seq chemistry.

Development of the scalability of the method continued with adaptation of the magnetic bead enrichment and the 10x Flex scRNA-seq assay from a tube based method into a 96 well plate format with reduced reaction volumes to enable high throughput analysis (Fig. 3a). This format also allowed for the integration of liquid handling automation to enable a single operator to process up to 96 samples simultaneously. This workflow was evaluated by analyzing matching cryopreserved whole blood (n=7) and density gradient isolated PBMC (n=1) samples using the automated plate format magnetic bead enrichment and FACS enrichment methods followed by 10x Flex scRNA-seq, resulting in 110,594 (Bead Enrichment) and 103,501 (FACS) individual recovered cells. Library quality was assessed by comparing genes per cell as recovered using the plate based magnetic bead enrichment (1743 ± 59 s.e.m.) and FACS (1762 ± 99 s.e.m.) methods (Fig. 3b). UMAPs of gene expression overlaid with major immune cell type labeling showed similar clustering between the two enrichment methods, and cell type proportions show high correlation (R=0.988, p<0.001), indicating no significant differences in PBMC frequency in either enrichment method (Fig. 3c). Differences in differentially expressed genes between donors were conserved in both enrichment methods, and the gene expression between enrichment methods are highly correlated (R>0.988) (Fig. 3d). The magnetic bead depletion in plate format produced high quality results comparable to FACS enrichment, enabling us to analyze samples with much higher throughput at significantly reduced cost.

**Figure 3:**
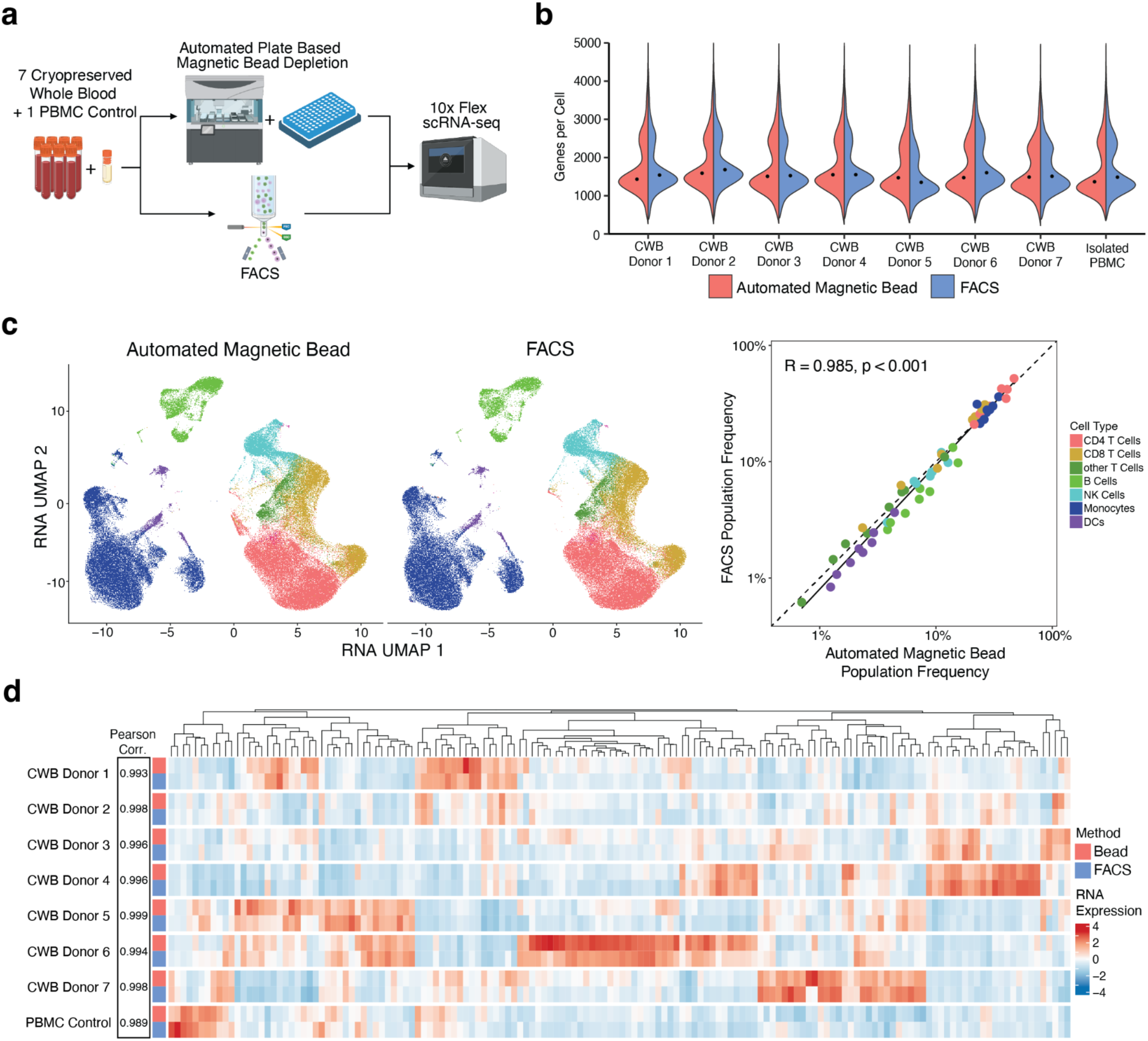
Scaling of cryopreserved whole blood sample throughput with multiplexed scRNA-seq and automated sample handling. **a.** Overview of experimental method for evaluating cryopreserved whole blood (n=7) and PBMC (n=1) donors in plate based workflow incorporating magnetic bead enrichment with automated liquid handling compared to FACS. **b.** Genes per cell recovered from individual donors automated magnetic bead enrichment (red) and FACS (blue) **c.** (Left) UMAP visualization clustered on gene expression data and overlaid with cell type labels comparing the automated magnetic bead enrichment and FACS. (Right) Correlation of cell type population frequencies between enrichment methods (Dashed line equivalent to R=1, solid line equivalent to regression line, R=0.988, p<0.001). **d.** Relative gene expression levels for the highest expressing genes per cell type for each enrichment method. High correlation was observed between donor replicates indicating consistent gene detection between methods (R>0.993).

Utilizing multiplexed scRNA-seq, plate based magnetic bead enrichment, and automated sample handling enabled us to develop a high throughput, cost efficient workflow for gene expression analysis on fixed cells from cryopreserved whole blood (Fig. 4a). To verify this workflow, 60 cryopreserved whole blood samples and 1 PBMC control were processed simultaneously, generating a dataset containing scRNA-seq data from 429,366 single cells (Fig. 4b). Samples were divided into 4 pools of 16 samples, where each pool was thawed by a separate operator. Within each pool, 3 replicate cryopreserved whole blood tubes from 5 individual donors were analyzed in order to evaluate consistency within and between pools. A single density gradient isolated PBMC sample was divided into 4 wells and added to each sample pool as a bridging control to further assess technical variance.

**Figure 4:**
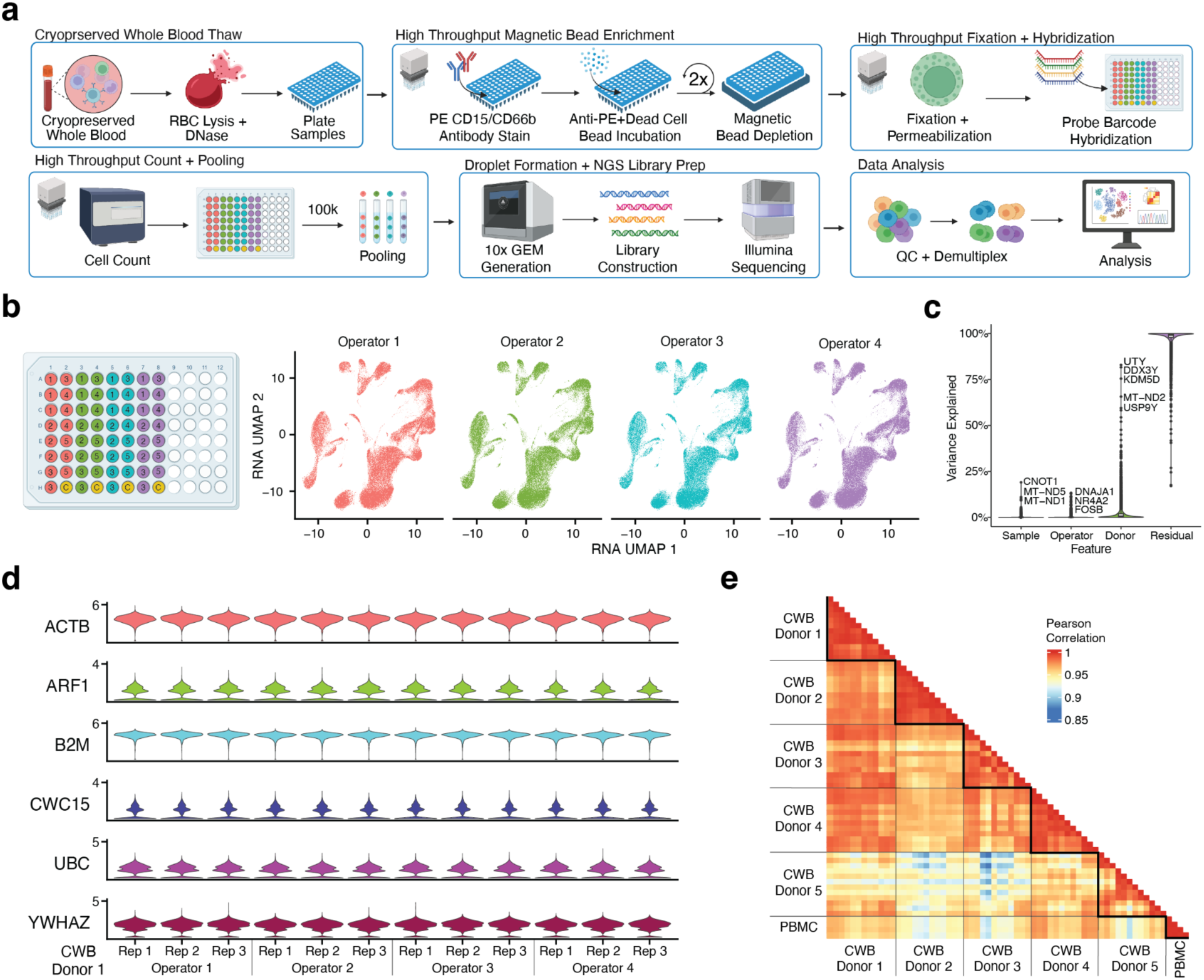
Verification of high throughput workflow with 60 cryopreserved whole blood samples through automated magnetic bead enrichment and scRNA-seq preparation. **a.** Overview of complete cryopreserved whole blood processing workflow. **b.** (Left) Layout of batch structure including 3 technical replicates of 5 donors across 4 thaw operators, and 1 PBMC batch control across all 4 pools. (Right) UMAP visualizations of single cell gene expression across each pool. **d.** Variance decomposition analysis evaluating contributions of sample technical replicates, operator, and donor to total variance of individual genes. The vast majority of variance in gene expression is not attributed to experimental variables **d.** Gene expression of 6 housekeeping genes across technical replicates from 1 donor, demonstrating high consistency between thaw operators and sample tubes (CVs for all genes <5%). **e.** Average gene expression correlation between each sample, showing high consistency between technical replicates (R>0.96).

Individual cryopreserved whole blood replicates exhibited consistent gene expression results with high correlation measured between replicates of the same donor (R>0.980). UMAPs of gene expression results between operators also display highly similar clustering (Fig. 4b, Supplement Figure 3). Variance decomposition analysis was performed to evaluate contributions of study variables to total variance of individual genes (Fig. 4c). Sample technical replicate and operator features were found to have low impact on the variance in gene expression (<10%). While some variance can be attributed to sex-linked genes from male and female donors (2 male, 6 female), the majority of gene expression variance can be attributed to biological factors, rather than deviations introduced by the workflow^21^. Within a single donor, gene expression from six housekeeping genes^22^ (ACTB, ARF1, B2M, CWC15, UBC, YWAHZ) across the four operators displayed consistent gene expression (coefficient of variation<5%), again demonstrating the methods reproducibility across replicates (Fig. 4d). Within the entire batch of samples, average gene expression was highly correlated between technical replicates (R>0.970), while conserving differences between donors (Fig. 4e). The generation of high quality and robust technical replicate data using this workflow demonstrates both the consistency of cryopreserved whole blood samples, even with handling by different operators, and the feasibility of using automation to increase sample throughput and reduce cost.

### Enabling high throughput functional assays on cryopreserved whole blood

A crucial component to immunological studies involving PBMCs is the ability to perturb collected cells and observe their response to stimuli with a variety of readouts. In addition to single cell readouts, we evaluated if samples preserved with the described cryopreserved whole blood method are compatible with functional assays at scale using well characterized stimulations that target different immune cell types. Cryopreserved whole blood (n=3) and density gradient isolated PBMC (n=1) samples were thawed and enriched for PBMCs using the automated magnetic bead depletion described previously. The transcriptional response of each sample to three stimulation conditions was measured, including phorbol 12-myristate 13-acetate (PMA) with ionomycin, resiquimod (R848), and CD3/CD28 T cell activation beads against unstimulated cells (Fig. 5a). After a four hour incubation period, cells were immediately fixed to preserve the transcriptional state, and then analyzed using 10x Flex scRNA-seq.

**Figure 5:**
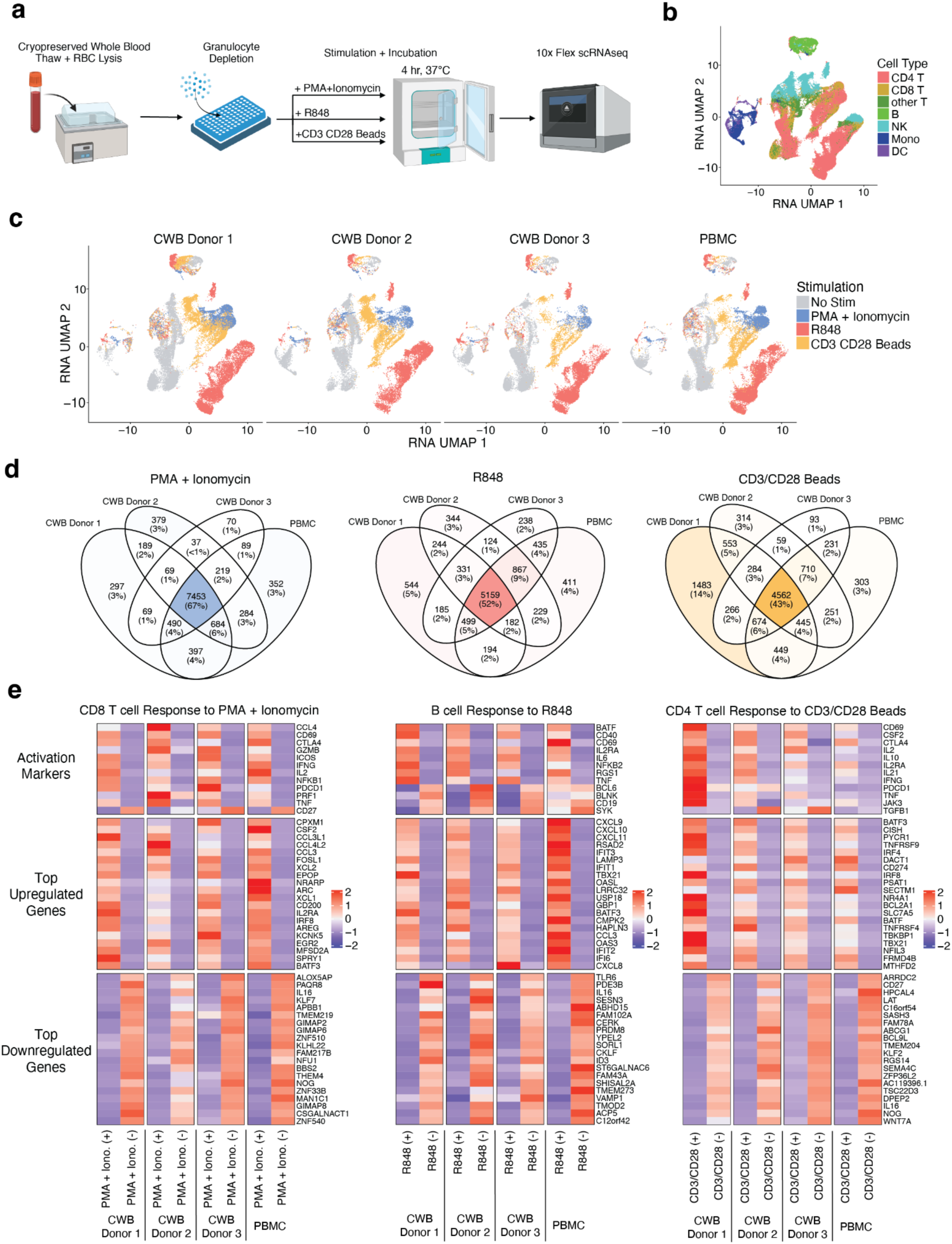
Functional assays using cryopreserved whole blood and multiplexed scRNA-seq. **a.** Overview of experimental method for stimulation of cryopreserved whole blood samples with PMA + ionomycin, R848, and CD3/CD28 T cell activation beads. **b.** UMAP visualization of gene expression from stimulated cryopreserved whole blood samples overlaid with major cells types. Separate islands of the same cell type indicate divergent gene expression from different stimulation conditions. **c.** UMAP visualization of gene expression from stimulated cryopreserved whole blood samples overlaid with specific stimulation conditions. Cells from each donor cluster together in response to stimulation, demonstrating preservation of cellular response in cryopreserved whole blood. **c.** Venn diagrams showing overlapping differentially expressed genes (DEGs) from individual donors in response to each stimulation condition. The majority of DEGS are common in all donors. **e.** Relative gene expression of known activation markers and unsupervised DEGs from all donors in response to each stimulation condition. Similar activation patterns are seen in all donors.

UMAP projections of the gene expression data from these functional assays reveal specific cell types grouping in multiple clusters, representing distinct responses to stimulation (Fig. 5b). Cells from all donors also clustered together in response to each specific stimulation, demonstrating the preservation of cellular response in cryopreserved whole blood compared to density gradient isolated PBMC (Fig. 5c, Supplement Fig. 4). Analysis of each stimulation condition against matched unstimulated cells also revealed a high degree of concordance of differentially expressed genes (DEGs) for all donors irrespective of cryopreservation method (67% PMA + Iono., 52% R848, 42% CD3/CD28 Beads) (Fig. 5d). Some DEGs remain specific to individual donors, including a strong DEG response signature to CD3/CD28 Beads occurring in one cryopreserved whole blood donor, indicative of the maintenance of biological differences between donors in the whole blood cryopreservation method.

A deeper look into specific activation marker genes reveals classical cell type specific responses in all donors, including CD8 T cell response to PMA and Ionomycin, B cell response to R848, and CD4 T cell response to CD3/CD28 activation beads (Fig. 5E) ^23,24^. Unbiased analysis of the highest upregulated and downregulated genes also show comparable patterns between all cryopreserved whole blood and density gradient isolated PBMC donors. Confirmation of these expected gene signatures and cell type specific responses enables the use of this cryopreserved whole blood method for scalable functional assays, and further validates the importance of this method for large scale studies or settings where PBMC processing is operationally unfeasible.

## Discussion

While technology to interrogate the immune system continues to rapidly innovate, sample preservation methods have had limited improvements to enable large scale clinical investigations and remain a barrier to studying underserved communities remote from research facilities. Density gradient isolation and cryopreservation of PBMCs have been the gold standard for longitudinal immune cell studies for decades, but the drawbacks to this method, including laboratory requirements and processing delays, have persisted. Alternative methods of preserving PBMCs using fixation do address these issues, but only allow for analysis of the samples at the time of blood draw and prevent use in functional assays. We developed a simplified workflow for cryopreservation of whole blood that enables immediate freezing of whole blood draws, allows for generation of high quality data from sequencing readouts, and facilitates functional assays on preserved cells. The CryoSCAPE method enables clinical studies using blood draws to collect and preserve samples immediately, only requiring cold storage (-80°C freezer or dry ice) with minimal training.

Similar whole blood cryopreservation methods have been described previously and demonstrated this whole blood cryopreservation method is viable for flow cytometry and scRNA-seq analysis, but without faithfully maintaining cell proportions or demonstration with large scale studies. We presented a method for whole blood cryopreservation that is compatible with deep molecular profiling including highly multiplexed fixed RNA-seq, CITE-seq, and ATAC-seq. Additionally, the ability to apply RNA-seq and CITE-seq on fixed cells from this cryopreservation method increases the throughput and reduces the cost of running these assays. Other methodologies using DMSO whole blood cryopreservation have indicated loss of monocytes compared to matched PBMC samples due to use of red blood cell depletion beads that remove platelets bound to monocytes, and require high amounts of EDTA to be supplemented to prevent this effect^14^. The use of an RBC lysis buffer in this method to remove red blood cells has no impact on cell type frequency or transcriptional readouts.

We also demonstrated that this method of cryopreservation is compatible with high throughput studies utilizing liquid handling automation, including functional studies incorporating large numbers of conditions or samples. Use of fixation prior to scRNA-seq can also allow for very short and long time scale functional assays, further improving upon other cryopreservation methods that fix the cells at the time of collection, or have only been demonstrated to work with unfixed sequencing assays. In conclusion, our study presents a significant advancement in sample preservation methods and post-thaw processing for large-scale clinical investigations focusing on peripheral blood mononuclear cells (PBMCs). Our proposed method of cryopreserving whole blood addresses clinical sample collection challenges by enabling immediate freezing of blood samples, thus streamlining workflow and facilitating high-quality data generation across various sequencing readout assays. Moreover, our approach is not only compatible with functional assays but also demonstrates reproducibility and correlation across multiple sequencing pools and sample thaw operators.

### Conclusions

Incorporation of this cryopreservation methodology and high throughput automation into clinical trials has the potential to transform our understanding of immunology. Simplified blood collection on the clinical side opens up understudied geographic, economic, racial and ethnic populations that previously had poor representation in studies due to the inability to reach clinical centers that could process and cryopreserve PBMCs. Scaling of sample throughput using automation and targeted multi-omic readouts drastically lowers the cost per sample and allows research centers to increase the power of their studies. By offering a simplified and efficient alternative to conventional methods, CryoSCAPE opens doors for enhanced research capabilities in immunology and beyond, promising greater accessibility and reliability in clinical studies utilizing blood samples.

## Methods

### Sample Collection

Whole blood draws were collected with written informed consent under the supervision of an Institutional Review Board (IRB) Protocol. Whole blood sourced from healthy subjects were collected by BloodWorks NorthWest (Seattle, WA) and the Benaroya Research Institute (Seattle, WA) through protocols approved by the relevant institutional review boards (Supplement Table 1). Blood samples were drawn into blood collection tubes with sodium heparin, and delivered to the Allen Institute within 4 hours of blood draw and processed immediately.

### Whole Blood Cryopreservation

Cryopreserved whole blood aliquots were generated by combining 2 mLs of fresh whole blood with 2 mLs of room temperature 15% DMSO freezing media containing CryoStor10 (Biolife Solutions) supplemented with additional DMSO (Sigma). Whole blood and freezing media was combined in a 5 mL cryotube, resulting in a final freezing concentration of 7.5% DMSO. Cryotubes were capped and inverted 10 times to fully mix the blood and freezing media. Samples were then transferred to a room temperature CoolCell Freezing Container (Corning) and stored in a -80°C freezer for 24 hours to slow freeze at a rate of 1°C per hour. Frozen whole blood aliquots were then stored in vapor phase liquid nitrogen.

### PBMC Isolation and Cryopreservation

Peripheral blood mononuclear cells (PBMCs) were isolated from whole blood by Ficoll density gradient (GE Healthcare) centrifugation using SepMate 50mL Isolation tubes (STEMCELL Technologies) according to the manufacturer’s protocol. The PBMC fraction was isolated and resuspended in CryoStor CS10 (Biolife Solutions) at a concentration of 5 million cells per mL. PBMC suspensions were distributed in 1 mL cell aliquots into 1.5 mL cryotubes, transferred to a Corning CoolCell Freezing Container, and stored in a -80°C freezer for 24 hours. Frozen PBMC aliquots were then stored in vapor phase liquid nitrogen.

### Cryopreserved Whole Blood Thaw

Cryopreserved whole blood samples were thawed in a 37°C water bath, transferred into 30 mL of pre-warmed AIM V media (37°C) (GIBCO), and centrifuged for 10 minutes at 400g (4°C) in a swinging bucket rotor. Thawed cells were washed with 30 mL of cold AIM V media (4°C) and centrifuged for 10 minutes at 400g (4°C) in a swinging bucket rotor. To remove red blood cells (RBC), the thawed samples were resuspended in 10 mL of cold RBC Lysis Buffer (BioLegend) for 15 minutes (4°C). 20 mL of AIM V media (4°C) was added to quench the lysis reaction, and then cells were centrifuged for 10 minutes at 400g (4°C). To prevent clumping, thawed cells were resuspended in 200 µL of 0.2 mg/mL DNase I Solution (STEMCELL Technologies), incubated at room temperature for 15 minutes. The cells were then washed with 30 mL of cold AIM V media (4°C) and centrifuged for 10 minutes at 400g (4°C). Individual samples were resuspended in 250 µL of Dulbecco’s phosphate-buffered saline (DPBS) (Costar) for cell staining for FACS, or in 250 µL of Mojosort Isolation Buffer (BioLegend) for magnetic bead enrichment.

### Cryopreserved PBMC Thaw

Cryopreserved PBMC samples were thawed in a 37°C water bath, transferred into 30 mL of pre-warmed AIM V media (37°C) (GIBCO), and centrifuged for 10 minutes at 400g (4°C). Thawed cells were resuspended in 5 mL of cold AIM V (4°C), and 30 µL was removed for counting. The counting fraction was stained 1:1 with 30 uL of AO/PI solution and counted on a Nexcelom Cellaca MX cell counter (Supplement Note). Thawed cells were then washed with 25 mL of cold AIM V media (4°C) and centrifuged for 10 minutes at 400g (4°C). Individual samples were resuspended in Dulbecco’s phosphate-buffered saline (DPBS) (Costar) at a concentration of 10 million cells per mL for cell staining for FACS, or in 250 µL of Mojosort Isolation Buffer (BioLegend) for granulocyte and dead cell bead enrichment.

### FACS and Flow Cytometry Analysis

Fluorescence activated cell sorting (FACS) and flow cytometry analysis were performed using a 3 marker panel including FVS510 Fixable Viability Stain (BD Biosciences), CD45 FITC (Clone 2D1, BioLegend), and CD15 PE (Clone HI98, BioLegend). Samples were stained 1:500 with FVS510 viability stain in 100 uL total volume for 20 minutes (4°C), and washed once with Mojosort Isolation Buffer (BioLegend). Samples were then stained with 3 uL each of CD15 PE and CD45 FITC antibodies in 100 uL total volume for 30 minutes (4°C), and washed twice and resuspended in Mojosort Isolation Buffer. Viable PBMCs were sorted with a BD FACSAria cell sorter, gated on FVS510 viability negative, CD15 PE negative, and CD45 FITC positive cells (Supplement Fig. 1a). Bead enriched cells were analyzed using a Cytek Aurora spectral cytometer (Supplement Note). Compensation and unmixing controls for each respective instrument were generated from healthy PBMC samples. Flow Cytometry Standard (FCS) files from the BD FACSArea cell sorter and the Cytek Aurora spectral cytometer were exported in FCS 3.1 format. Manual gating analysis for figures was generated using FlowJo (BD Biosciences, v10.10.0).

### Magnetic Bead Enrichment

Samples were stained with 5 uL CD15 PE (Clone HI98, BioLegend) and 5uL CD66b PE (Clone 6/40c) in a total volume of 100 uL for 15 minutes (4°C). Stained cells were washed with 100 uL of 1X Mojosort Isolation Buffer (Biolegend), centrifuged at 400g for 5 min (4°C), resuspended in 100uL of 1X Mojosort Isolation Buffer. Samples were then stained with 5 uL of Mojosort Human Dead Cell Removal Beads (BioLegend) for one round bead depletion, or 5 uL of Mojosort Human Dead Cell Removal Beads and 5 uL of Mojosort Human anti-PE Nanobeads (BioLegend) for two round bead depletion. Samples were incubated with magnetic beads for 15 minutes (4°C). Samples stained in tubes were washed with 3 mL of Mojosort Isolation Buffer, moved to a 5 mL tube magnet (Biolegend), and incubated for 3 minutes. Samples stained in plate format were washed with 150 uL of Mojosort Isolation Buffer, moved to a plate magnet adapter (Alpaqua), and incubated for 3 minutes. After transfering the supernatant to a new tube, the remaining beads were washed and magnetically depleted a second time. For samples undergoing two round depletion, the enriched cells were centrifuged at 400g for 5 min (4°C), stained a second time with 5 uL of Mojosort Human Dead Cell Removal Beads and 5 uL of Mojosort Human anti-PE Nanobeads, and magnetically enriched on the plate magnet two additional times. Enriched cells were centrifuged at 400g for 5 min (4°C) and resuspended in 280 uL of Dulbecco’s phosphate-buffered saline (DPBS). 30 uL of each sample was stained 1:1 with 30 uL of AO/PI solution and counted on a Nexcelom Cellaca MX cell counter (Supplement Note). Plate format bead isolation and cell counting pipetting steps completed using a Tecan Fluent 1080 liquid handling automation system (Supplement Note).

### 10x Chromium Flex scRNA-seq and scCITE-seq

PBMC enriched samples were processed according to the 10x Genomics protocol for Fixation of Cells and Nuclei (CG000478, Rev C). Volumes were scaled from 1 mL to 250 uL for plate-based sample preparation and handling. Prior to fixation, samples were stained using BioLegend’s TotalSeq™-C Human Universal Cocktail(Cat. No 399905) according to the TotalSeq™-B or -C with 10x Feature Barcoding Technology protocol. Following extracellular protein staining, samples were fixed in a final concentration of 4% Paraformaldehyde for 1 hour (25°C). Fixed samples were quenched and subsequently washed with an additional 200 uL of PBS + 0.02% BSA (Costar). Samples were barcoded for multiplexing using 10X Genomics Fixed RNA Feature Barcode Multiplexing Kit (PN-1000628). Probe hybridization was completed according to the 10x Genomics protocol for Multiplexed samples with Feature Barcode technology for Protein using Barcode Oligo Capture (CG000673, Rev A). Up to 2 million fixed cells were hybridized per sample. After 16-24 hours of probe hybridization incubation (42°C), samples were diluted in Post-Hyb Wash buffer and 11 µL was removed for counting. The counting fraction was stained 1:5 with 44 uL of PI solution and counted on a Nexcelom Cellaca MX cell counter (Supplement Note). Samples were then pooled at equivalent cell concentrations, incubated for 3 rounds of 10 minutes at 42°C, washed in Post-Hyb wash buffer, and strained using 30 uM strainers (Miltenyi). The single cell suspension pool was loaded onto Chip Q for GEM generation at an overloaded concentration of 400,000 cells. Final scRNAseq libraries were sequenced using a NovaSeq X 25B PE300 or NovaSeq S4 PE100 flow cell, targeting a read depth of 10,000 reads per cell.

### 10x Chromium 3’ v3.1 scRNA-seq

PBMC enriched samples were processed with the 10X Genomics Chromium Next Gem Single Cell 3’ kit (v3.1) using a previously published protocol^25^ <u>(Genge et al. 2021)</u>. Samples were stained with cell hashing antibodies and combined into pools of 12 samples. Pooled samples were then added to 10x Chromium Controller Chip G in replicate at a concentration of ∼64,000 cells per well across 12 wells. Generated libraries were sequenced on Illumina Novaseq using an S4 flowcell with 45k reads per cell for RNA libraries and 3k reads per cell for HTO libraries.

### Bulk ATAC-seq

PBMC enriched samples were first permeabilized using 0.1% digitonin. Tagmentation reactions using Tagment DNA TDE1 Enzyme and Buffer Kit (Illumina, PN 20034198) were carried out on 50,000 cells incubated for one hour at 37°C. DNA products were purified using a DNA Clean-Up buffer containing 0.1% SDS, 16 mM EDTA, 160 ug Proteinase K (ThermoFisher, PN EO0491), 23 ug RNase A (ThermoFisher, PN EN0531) and incubated at 37°C overnight, followed by a 1.8X AMPure XP bead selection (Beckman Coulter). DNA products were barcoded with a unique i7 primer and an universal i5 primer^26,27^ and purified using a 0.4X right-sided and 1.2X left-sided AMPure XP bead clean up. Libraries were sequenced using a NextSeq2000 P2 PE50 flow cell, with a sequencing depth of 10,000,000 reads per library.

### PBMC Stimulation

PBMC enriched samples were resuspended in complete RPMI (Roswell Park Memorial Institute 1640 media + 10% Fetal Bovine Serum + 1% Penicillin Streptomycin, Gibco) and plated in a 96 well U bottom plate at a concentration of 500,000 cells per well. Cells were rested for 1 hour at 37°C, and then exposed to one of four stimulation conditions: no stimulation, phorbol 12-myristate 13-acetate (PMA) (125 ng/mL, Fisher Scientific) with ionomycin (2.5 μg/mL, Fisher Scientific), resiquimod (R848) (500 ng/mL, Invivogen), or CD3/CD28 T cell activation beads (0.5 beads/cell, Gibco). Cells were incubated for 4 hours at 37°C, then resuspended in 4% PFA for fixation and subsequent 10x Flex scRNA-seq analysis.

### scRNA-seq and scCITE-seq Analysis

FASTQ files were used in the Cell Ranger (10x Genomics, v7.1.0) alignment function against the human reference annotation (Ensembl GRCh38). 10x Flex scRNA-seq and scCITE-seq matrices were generated using Cell Ranger Multi (10x Genomics, v7.1.0). 10x 3’ v3.1 scRNA-seq matrices were processed using Cell Ranger, as described in <u>Savage et al. 2021</u> ^8^ and <u>Swanson et al. 2021</u> ^28^. Cell doublets were identified in the data using Scrublet (v0.2.3) ^33^ Filtered cell by gene matrices were processed using Scrublet and saved to a comma separated value (csv) file with cell barcode, doublet score and doublet classification. Filtered cell by gene matrices were read into Seurat (Satija Lab, v5.1.0) with the Scrublet results to remove doublets from the analysis. Datasets were further filtered to remove cells with greater than 5% of mitochondrial gene expression.

To analyze scRNA-seq data, the filtered matrices were normalized (NormalizeData function, default settings), scaled (ScaleData function, defaults), and analyzed with principal component analysis (PCA) (RunPCA function, defaults). UMAPs (RunUMAP, dims = 1:20) and clustering analysis (FindClusters, resolution = 0.5, FindNeighbors, dims = 1:20) were also completed using the Seurat package. For label transfer, SCTransform was used to normalize the data in order to match the Seurat Multimodal Reference Dataset for PBMCs (available from the Satija lab at https://atlas.fredhutch.org/data/nygc/multimodal/pbmc_multimodal.h5seurat, Hao et al., 2020). Label transfer was then performed using the Seurat functions FindTransferAnchors and TransferData as described in the Seurat v4 Vignettes. Reference labels were mapped in the ‘celltype.l1’ categories onto the filtered datasets. Cells labeled as celltype “other” was dropped from further analysis. For ADT counts, the filtered matrices were normalized (NormalizeData function, normalization.method=’CLR’, margin = 2), scaled (ScaleData function, defaults), and analyzed with principal component analysis (PCA) (RunPCA function, defaults). UMAPs (RunUMAP, dims = 1:20) and clustering analysis (FindClusters, resolution = 0.5, FindNeighbors, dims = 1:20) were also completed using the Seurat package. Variance decomposition was performed using the PALMO package as described in Vasaiker et al. 2023 [21]. Processed Seurat objects for all data types were saved as .rds files for plotting.

### Bulk ATAC-seq Analysis

Bulk ATAC-Seq samples were run through an internal bulk ATAC-Seq analysis pipeline in which each read pair is aligned to hg38 using bowtie2 (v2.4.5, available at https://github.com/BenLangmead/bowtie2). Results are written out to a SAM file and used to create a corresponding BAM file using samtools and a sorted BED file using bedtools for downstream analysis. To compare fragment length distributions between donors and sample conditions, the BED files are used to calculate fragment widths and fraction of total reads. Sorted BED files are randomly downsampled to 14 million reads and converted to BedGraph files as an intermediate to convert to BigWig files to complete genome track comparisons. The downsampled BED files are also used to examine transcription start site footprint position and coverage overlap using GRanges and IRanges relative to 2kb upstream and downstream of the TSS. Genome track plots were generated using Integrative Genomics Viewer^34^.

### Statistical Analysis

Pearson correlation coefficients and p-values for cell frequency, gene expression, and protein expression comparisons across all figures were calculated using the *cor* and *cor.test* functions in the *stats* base package (v3.6.2) in the R statistical programing language (v4.3.3).

## Abbreviations

ADT: Antibody derived tag
CryoSCAPE: Cryopreservation for Scalable Cellular And Protein Exploration
CWB: Cryopreserved whole blood
DC: Dendritic Cells
DEG: Differentially expressed gene
DMSO: Dimethyl sulfoxide
DPBS: Dulbecco’s phosphate-buffered saline
EDTA: Ethylenediaminetetraacetic acid
FACS: Fluorescence activated cell sorting
IRB: Institutional review board
NK: Natural killer
PBMC: Peripheral blood mononuclear cell
RBC: Red blood cell
TSS: Transcriptional start site
UMAP: Uniform manifold approximation and projection
UMI: Unique molecular identifier

## Declarations

### Ethics approval and consent to participate

Specimens purchased from Bloodworks NW were collected under BloodWorks WIRB Protocol #20141589. Specimens provided by Benaroya Research Institute were collected under Benaroya Research Institute IRB #IRB19-045. All sample collections were conducted by Bloodworks NW and Benaroya Research Institute under these IRB protocols, and all donors signed informed consent forms.

### Consent for publication

Not applicable.

### Data Availability

Raw 10x Flex scRNA-seq data has been deposited in SRA (accession number # SUB14488403). Raw 10x 3’ v3.1 scRNAseq data are deposited dbGaP for controlled access (dbGaP study accession #). Processed scRNA-seq (10x Flex and 10x 3’ v3.1), scCITE-seq, and bulk ATAC data has been deposited in GEO (accession number #). All graphical figures created using Biorender.com. Data processing notebooks are available via Zenodo (10.5281/zenodo.11479595). Analysis code is available on Github (https://github.com/aifimmunology/CWB_Paper).

### Authors’ contributions

A.H., C.P., N.K., P.W., J.G., M.W., and P.S. designed the experiments, analyzed the data and prepared the manuscript. P.G. processed ATAC-seq data and provided bioinformatics support. Z.T., C.G., and J.R., provided experimental, analysis, and editing support.

## Acknowledgments

The authors are grateful to the donors that provided biological material for this study. The authors would like to thank the members of the Allen Institute for Immunology, especially the Lab Operations team who helped facilitate this research, as well as the Human Immune System Explorer (HISE) software development team for their support. The authors also thank the Allen Institute founder, Paul G. Allen, for his vision, encouragement, and support.

