## Supplementary Material for "CryoSCAPE: Scalable Immune Profiling Using Cryopreserved Whole Blood for Multi-omic Single Cell and Functional Assays"



© Allen Institute for Immunology, All Rights Reserved (confidential)

**Supplement Table 1: Sample Donor Key**

| Figure | Figure Donor ID | Sample Type | AIFI Donor ID | Sex | Age | Source |
|---|---|---|---|---|---|---|
| 1 | Donor 1 | Matched CWB/PMBC | BL05111 | Male | 27 | Bloodworks NW |
| | Donor 2 | Matched CWB/PMBC | BL05113 | Male | 29 | Bloodworks NW |
| | Donor 3 | Matched CWB/PMBC | BL05402 | Female | 56 | Bloodworks NW |
| 2A, 2B, 2C | Donor 1 | PBMC | PB00607 | Male | 32 | Benaroya Research Institute (BRI) |
| | Donor 2 | PBMC | PB00609 | Male | 31 | Benaroya Research Institute (BRI) |
| | Donor 3 | PBMC | PB00610 | Female | 36 | Benaroya Research Institute (BRI) |
| 2D | CWB Donor 1 | CWB | BL05012 | Female | 40 | Bloodworks NW |
| | CWB Donor 2 | CWB | BL05044 | Female | 68 | Bloodworks NW |
| | CWB Donor 3 | CWB | BL05045 | Female | 63 | Bloodworks NW |
| | PBMC | PBMC | PB04745 | Female | 55 | Bloodworks NW |
| 2E | CWB Donor 1 | CWB | BL05731 | Female | 40 | Bloodworks NW |
| | PBMC | PBMC | PB04745 | Female | 55 | Bloodworks NW |
| 3 | CWB Donor 1 | CWB | BL05012 | Female | 40 | Bloodworks NW |
| | CWB Donor 2 | CWB | BL05013 | Male | 38 | Bloodworks NW |
| | CWB Donor 3 | CWB | BL05044 | Female | 68 | Bloodworks NW |
| | CWB Donor 4 | CWB | BL05045 | Female | 63 | Bloodworks NW |
| | CWB Donor 5 | CWB | BL05729 | Female | 72 | Bloodworks NW |
| | CWB Donor 6 | CWB | BL05730 | Female | 64 | Bloodworks NW |
| | CWB Donor 7 | CWB | BL05731 | Female | 40 | Bloodworks NW |
| | PBMC | PBMC | PB02183 | Male | 28 | Bloodworks NW |
| 4 | CWB Donor 1 | CWB | BL05013 | Male | 38 | Bloodworks NW |
| | CWB Donor 2 | CWB | BL05045 | Female | 63 | Bloodworks NW |
| | CWB Donor 3 | CWB | BL05113 | Male | 29 | Bloodworks NW |
| | CWB Donor 4 | CWB | BL05759 | Male | 45 | Bloodworks NW |
| | CWB Donor 5 | CWB | BL05760 | Female | 65 | Bloodworks NW |
| | PBMC | PBMC | PB02183 | Male | 28 | Bloodworks NW |
| 5 | CWB Donor 1 | CWB | BL05012 | Female | 40 | Bloodworks NW |
| | CWB Donor 2 | CWB | BL05111 | Male | 27 | Bloodworks NW |
| | CWB Donor 3 | CWB | BL05731 | Female | 40 | Bloodworks NW |
| | PBMC | PBMC | PB02183 | Male | 28 | Bloodworks NW |


© Allen Institute for Immunology, All Rights Reserved (confidential)

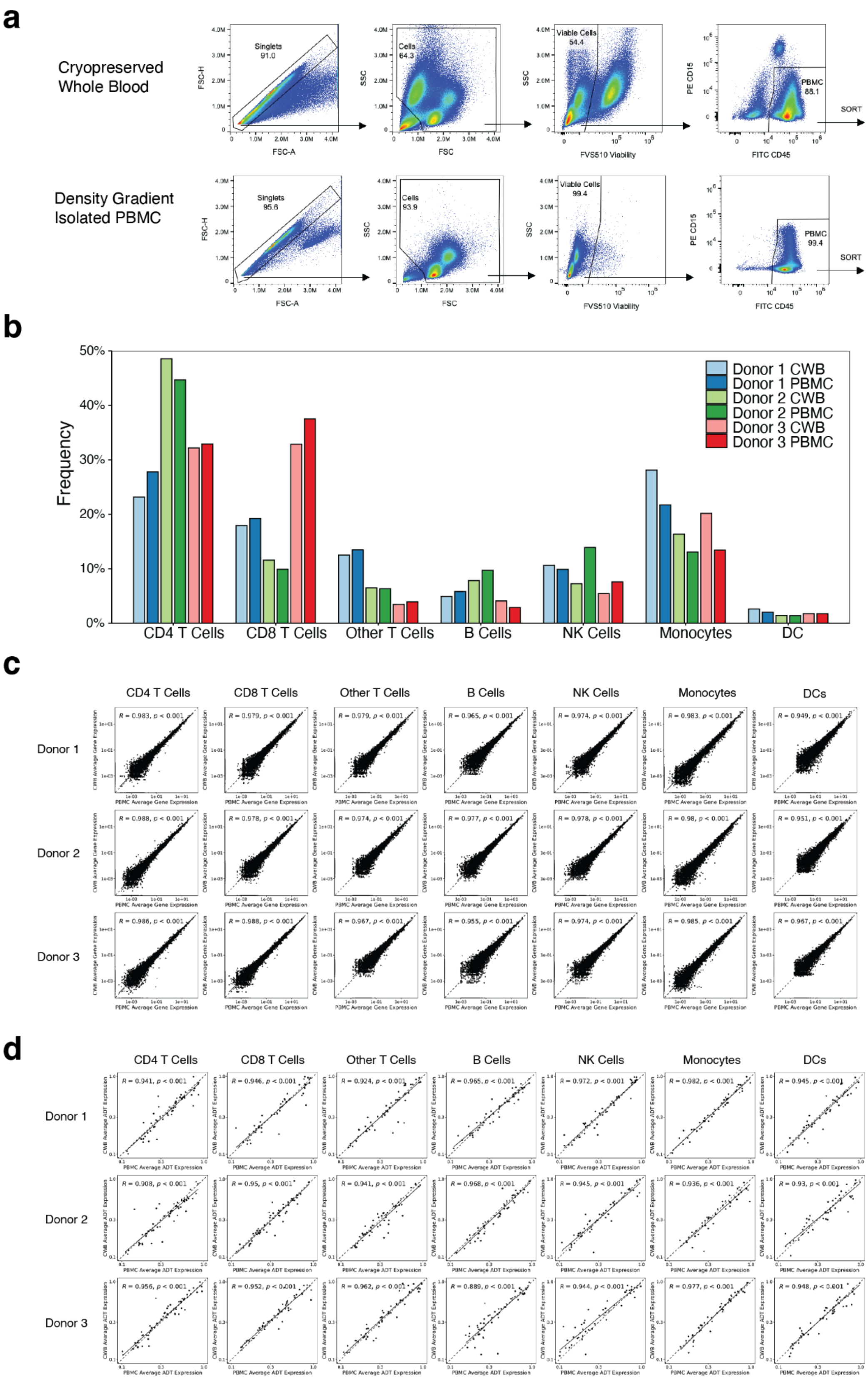


16



© Allen Institute for Immunology, All Rights Reserved (confidential)

**Supplement Figure 1: Whole blood cryopreservation in comparison to density gradient isolated PBMC cryopreservation for clinical site processing and multi-modal sequencing assays. a.** FACS gating schemes for cryopreserved whole blood and density gradient isolated PBMC samples. **b.** Cell type frequencies for matched cryopreserved whole blood and density gradient isolated PBMC samples. **c.** Gene expression correlation plots for all 10x Flex scRNA-seq targets between cryopreserved whole blood and density gradient isolated PBMC samples. **d.** Protein expression correlation plots for all sCITE-seq ADT targets between cryopreserved whole blood and density gradient isolated PBMC samples.


© Allen Institute for Immunology, All Rights Reserved (confidential)

**Supplement Table 2: Bulk ATAC-Seq Summary Metrics Table**

| Donor ID and Condition | Read Count | % Reads in Peaks | % Reads in TSS | % Mitochondrial Reads |
|---|---|---|---|---|
| Donor 1 (PBMC) | 18,546,680 | 19.00 | 8.00 | 1.22 |
| Donor 1 (CWB) | 19,933,577 | 20.00 | 8.00 | 1.20 |
| Donor 2 (PBMC) | 14,814,264 | 21.00 | 10.00 | 2.00 |
| Donor 2 (CWB) | 14,031,861 | 19.00 | 8.00 | 1.35 |
| Donor 3 (PBMC) | 19,258,569 | 21.00 | 10.00 | 1.59 |
| Donor 3 (CWB) | 17,798,421 | 19.00 | 8.00 | 1.12 |


© Allen Institute for Immunology, All Rights Reserved (confidential)

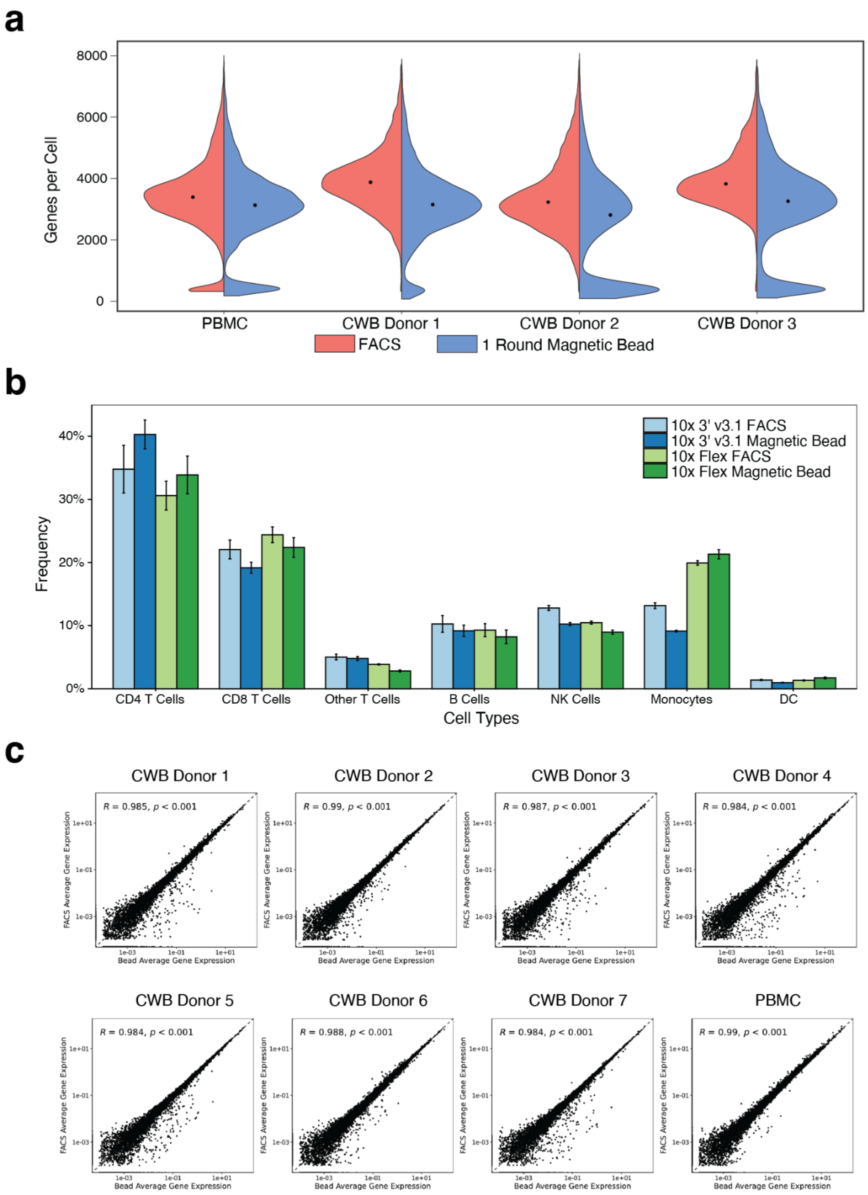


**Supplement Figure 2: Development 10x Flex scRNA-seq methodology as a high throughput method for single cell transcriptome analysis of cryopreserved whole blood.** **a.** Genes per cell recovered using the 10x 3' v3.1 chemistry with cryopreserved whole blood and density gradient isolated PBMC samples enriched using FACS (red) versus a single round of magnetic bead based enrichment (blue). **b.** Cell type frequencies for cryopreserved whole blood samples measured with the 10x 3' v3.1 (blue) or the 10x Flex (green) scRNA-seq chemistries,



© Allen Institute for Immunology, All Rights Reserved (confidential)

and enriched with either FACS or magnetic bead based enrichment. **c.** Gene expression correlation plots for all 10x Flex scRNA-seq targets between FACS and magnetic bead enriched cryopreserved whole blood samples.


© Allen Institute for Immunology, All Rights Reserved (confidential)

### Supplement Table 3: 10x Flex vs 10x 3’ v3.1 Cost Estimates for 64 Sample Batch Size

| Batch Metrics | 10x Flex scRNA-seq | 10 3’ v3.1 scRNA-seq |
|---|---|---|
| Total Samples | 64 | 64 |
| Total Cells Loaded per Well | 400,000 (1 well) | 11,651 (2 wells) |
| Estimated Singlets Captured per Well | 213,042 | 6,658 |
| Estimated Singlets Captured per Sample | 13,315 | 13,315 |
| Sequencing Reads per Cell | 10,000 | 20,000 |
| Total Sequencing Reads | 9,696,969,697 | 17,044,480,000 |
| 10x Reagent Cost | $26,588 | $216,480 |
| Sequencing Cost, NovaSeqX | $7,100 | $14,200 |
| Total Cost | $33,688 | $230,680 |
| Cost Per Sample | $526 | $3,604 |


© Allen Institute for Immunology, All Rights Reserved (confidential)

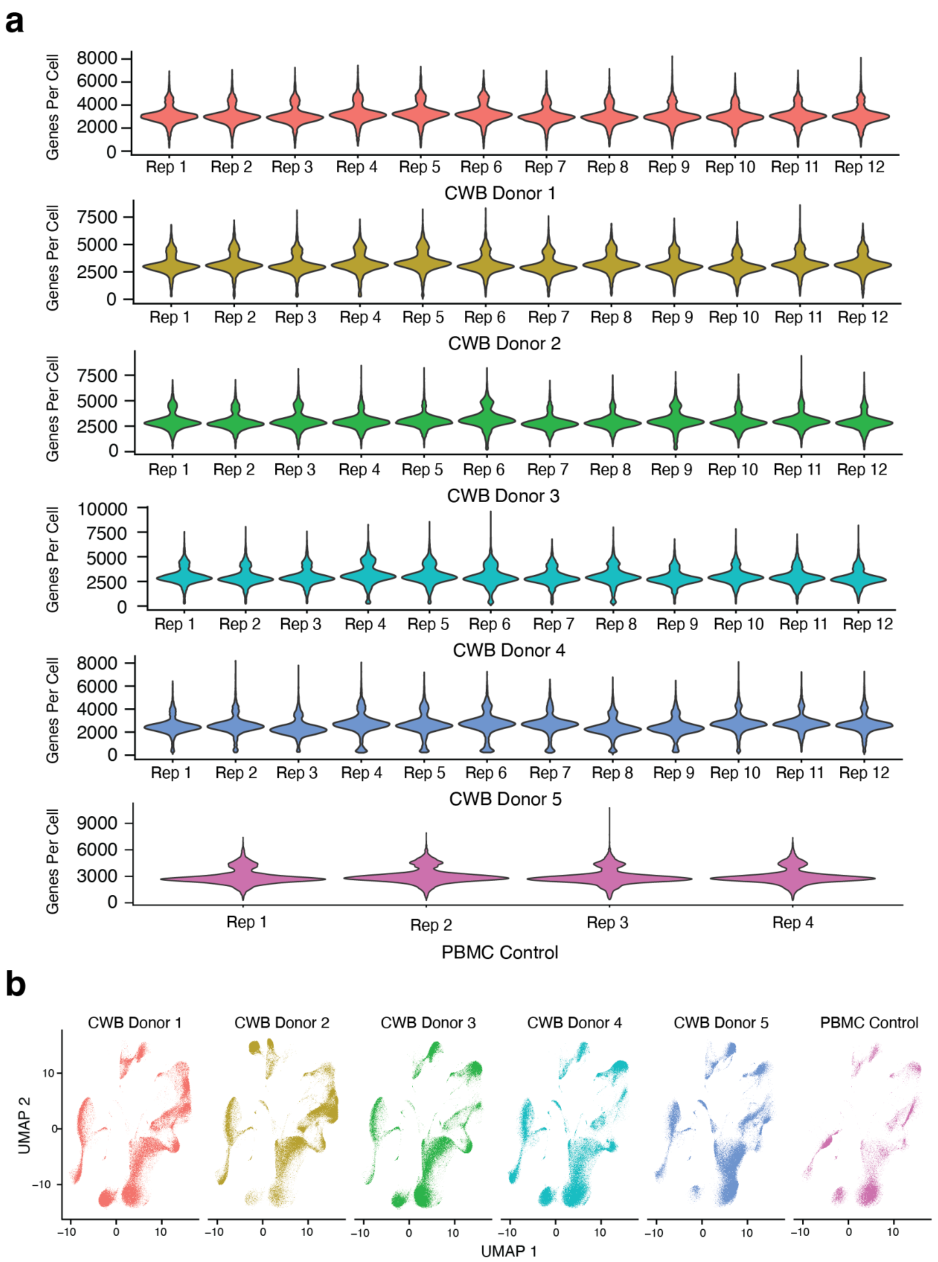


**Supplement Figure 3: Verification of high throughput workflow with 60 cryopreserved whole blood samples through automated magnetic bead enrichment and scRNA-seq preparation. a.** Genes per cell recovered using the 10x Flex chemistry with cryopreserved whole blood and density gradient isolated PBMC samples across all donors and replicates. **b.** UMAP visualizations of single cell gene expression across each donor.



© Allen Institute for Immunology, All Rights Reserved (confidential)

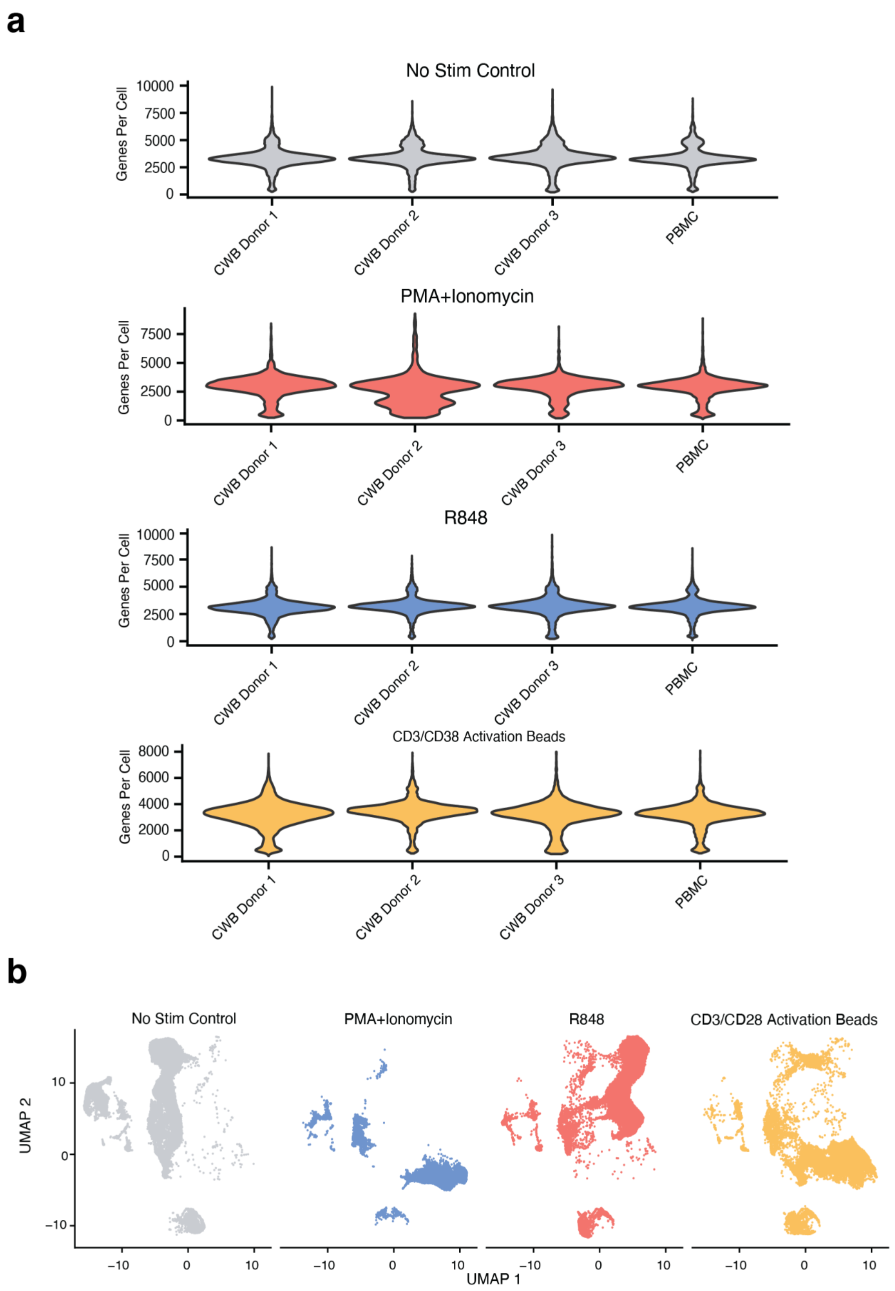


**Supplement Figure 4: Functional assays using cryopreserved whole blood and multiplexed scRNA-seq. a.** Genes per cell recovered using the 10x Flex chemistry with cryopreserved whole blood and density gradient isolated PBMC samples across all donors and stimulation conditions. **b.** UMAP visualizations of single cell gene expression across each stimulation condition.


© Allen Institute for Immunology, All Rights Reserved (confidential)

**Supplementary Note: Technical Information**

*Flow Cytometry and FACS*

Flow cytometry analysis was performed using a Cytek Aurora spectral flow cytometer. The analyzer was equipped with 5 lasers (ultra violet - 355 nm, violet - 405 nm, blue - 448 nm, yellow-green - 561 nm, red - 640 nm) and 64 fluorescent detectors. Instrument setup and performance tracking was performed daily using SpectroFlo QC beads (Cytek) using the Daily QC program. Experimental fluorescence gains were based on Cytek assay settings generated from Daily QC, while forward scatter (FSC-A) and side scatter (SSC-A) gains were optimized for healthy fixed PBMCs. The Aurora instrument was operated using Cytek SpectroFlo software version 2.2.0.4. All samples recorded on Medium flow rate (~30 uL/min) and event rate of ~5,000 events per second.

Fluorescent-activated cell sorting (FACS) was performed using a BD FACSAria Fusion cell sorter. The sorter was equipped with 4 lasers (violet - 405 nm, blue - 448 nm, yellow-green - 561 nm, red - 640 nm) and 14 fluorescent detectors. Instrument setup and performance tracking was performed daily using instrument specific Cytometer Setup and Tracking (CS&T) beads (BD) using the CS&T program. Experimental voltages were based on internal voltage optimization experiments with fixed healthy PBMCs. The FACSAria Fusion instrument was operated using BD FACSDiva software version 9.1. All samples sorted at a flow rate setting of 1 and event rate of ~5,000 events per second.

*Cell Counting*

Cell counting was performed using a Cellaca MX Cell Counter (Revvity) according to manufacturer specifications. Cells were stained 1:1 with Viastain Acridine Orange Propidium Iodine (AOPI) Staining Solution (Revvity) and 50uL plated into the counting wells of the 24 well



© Allen Institute for Immunology, All Rights Reserved (confidential)

counting plate (Revvity). The plate was loaded into the Cellaca MX and focus optimized per well. Live (AO) and dead cell (PI) populations were counted for viability and live cell calculations. Ratio of counted live cells to total counted cells was used to calculate viability.

For cell counts performed post-fixation, cells were stained with Viastain Propidium Iodine (PI) Staining solution (Revvity) 1:5 and 50uL plated into the counting wells of the 24 well counting plate (Revvity). The plate was loaded into the Cellaca MX and focus optimized per well. Total cell populations were counted to determine pooling normalization and GEM loading concentration.

*Automated Liquid Handling*

Automated liquid handling steps were completed using a Tecan Fluent 1080. The Tecan Fluent 1080 has a custom deck configuration including three CPAC 96 well plate heating/cooling units and 1 heating/cooling Thermoshake unit. A remote gripper arm (RGA) was used for labware transport, and flexible channel arm (FCA) or multiple channel arm (MCA) with extended volume adapter (EVA) attachment used for all liquid handling steps. Automated liquid handling scripts were developed and run using Tecan Fluent Control Build v2.7.19.54609. Deck layout and liquid handling scripts are available by request from the authors.


© Allen Institute for Immunology, All Rights Reserved (confidential)
